# Unveiling microglial heterogeneity from single-cell transcriptomics in neurodegenerative diseases

**DOI:** 10.1101/2024.08.23.609299

**Authors:** Alessandro Palma

## Abstract

Microglia are key players in maintaining brain homeostasis and responding to pathological conditions. Their multifaceted roles in health and disease have garnered significant attention in the context of neurodegeneration. In recent years, single-cell transcriptomic techniques have provided unprecedented insights into microglial heterogeneity, revealing distinct subpopulations and gene expression patterns associated with neuroprotection or neurotoxicity.

Here, the transcriptomic landscape of microglia has been dissected by leveraging human single-nuclei RNA sequencing datasets of neurodegenerative conditions, encompassing amyotrophic lateral sclerosis, frontotemporal dementia, Alzheimer’s disease, aging, and Parkinson’s disease. Results have led to the identification of distinct cell subpopulations, representative of the functional heterogeneity of the brain microglia. Moreover, distinct gene signatures and regulatory networks linked to inflammation and neurodegeneration have been identified.

Overall, the study provides an improved portrait of microglia in the context of neurodegenerative disorders, and it holds promise for developing a more targeted research aimed at modulating microglial function to mitigate disease progression and foster neuroprotection.

**Highlights:**

- Brain-resident microglia cells show a profound transcriptional heterogeneity
- Microglia are not subject to a macrophage-like polarization mechanism
- Neurodegenerative disorders share common transcriptional programs, yet retaining their own peculiarities
- Distinct gene regulatory networks underlie microglia heterogeneity and neurodegeneration-related dysfunctions

## Introduction

Microglia are the resident macrophages of the central nervous system (CNS), and play a crucial role in the developing brain, maintaining brain homeostasis, and responding to various pathological conditions (Li and Barres, 2018). These specialized phagocytes are now recognized as multifaceted actors in both health and disease, exerting numerous functions that are vital for the brain’s correct functioning. This include shaping the extracellular matrix surrounding synapses (Crapser et al., 2021), maintaining and regenerating the myelin sheath (Kent and Miron, 2024; Lloyd and Miron, 2019), and establishing crosstalk with other cells (Ronzano et al., 2021; Touil et al., 2023), among others.

In the context of neurodegenerative disorders, such as Alzheimer’s disease (AD), Parkinson’s disease (PD), amyotrophic lateral sclerosis (ALS), aging-related cognitive decline, and frontotemporal lobar dementia (FTLD), microglia have garnered significant attention due to their diverse functions and dynamic responses within the CNS (Gao et al., 2023; Hickman et al., 2018; Salter and Stevens, 2017). It is widely recognized that in AD, microglia are implicated in the clearance of amyloid-beta plaques (Hu et al., 2023), one of the hallmarks of the disease. However, dysfunctional microglia can contribute to neuroinflammation, exacerbating neuronal damage and cognitive decline (Leng and Edison, 2021). In PD, the involvement of microglia cells in disease pathogenesis is still under investigation, but their participation in neuroinflammation is increasingly recognized as a critical contributor to disease progression. Microglial activation, triggered by misfolded alpha-synuclein aggregates, leads to the release of pro-inflammatory cytokines and neurotoxic factors, contributing to dopaminergic neuron degeneration (Hickman et al., 2018; Joers et al., 2017). In ALS, microglia play a double role, exerting both neuroprotective and neurotoxic effects. While microglia initially attempt to clear misfolded proteins and damaged neurons, chronic activation can lead to the release of cytotoxic molecules, exacerbating motor neuron degeneration (Vahsen et al., 2021). Furthermore, in the context of neurodegeneration, including aging, microglia undergo phenotypic changes termed “microglial priming” (Niraula et al., 2017; Perry and Holmes, 2014). Primed microglia exhibit exaggerated inflammatory responses to stimuli and impaired resolution of inflammation, thus contributing to cognitive decline and increased vulnerability to neurodegenerative diseases. Finally, in FTLD, characterized by the selective degeneration of frontal and temporal lobes, microglia-mediated neuroinflammation is a prominent feature. Dysregulated microglial activation can lead to cytokine release, oxidative damage and mitochondrial dysfunction, thus contributing to synaptic dysfunction and neuronal loss in affected brain regions (Bright et al., 2019).

Understanding the complex roles of microglia in neurodegenerative disorders is essential for developing targeted therapeutic interventions aimed at modulating microglial function to mitigate disease progression and promote neuroprotection in affected individuals.

In recent years, the advent of single-cell transcriptomic techniques has revolutionized our understanding of cellular heterogeneity and dynamics. These cutting-edge methodologies have enabled researchers to dissect the transcriptomes of individual cells, providing unprecedented insights into the diversity and functional states of various cell types, including brain-resident cells. For instance, by profiling thousands of individual cells, studies have revealed previously unrecognized subpopulations within the microglial population (Masuda et al., 2020), each exhibiting distinct gene expression patterns and functional properties. By deciphering the transcriptional signatures associated with neuroprotective or neurotoxic microglial phenotypes, it could be possible to uncover potential therapeutic targets for modulating microglial responses in neurodegenerative disorders.

Here, an integration of distinct human single-nuclei RNA sequencing (snRNAseq) datasets of AD, AGING, ALS, FTLD, and PD patients, together with their cognate healthy controls, has been performed to dissect the transcriptomic landscape of microglia in both health and disease. The integrated analysis revealed important gene signatures and regulatory networks linked to inflammation, neurodegeneration, and aberrant cell behavior.

## Materials and methods

### Single cell RNA sequencing data processing

Publicly available single-nuclei RNA sequencing data have been downloaded from the Gene Expression Omnibus (GEO) repository under the following accession numbers: GSE243292, GSE219281, GSE174332. Parkinson’s disease data have been downloaded from https://doi.org/10.5281/zenodo.7886802 (Dehestani et al., 2023). Analyses have been conducted with the *Seurat v5* package (Hao et al., 2024) in the R environment. Each dataset has been separately pre-processed selecting only the microglial cell population whenever the dataset contained also other cell types. In particular, for the dataset with accession ID GSE219281, containing different brain cell populations, the microglia cell population has been selected using the following markers: “CX3CR1”,”C1QB”,”CD74”,”C3”. The dataset with accession ID GSE174332 already provided the annotation for the microglial cell population, which has been selected using the “*subset*” function. Data have been normalized, scaled and the top 2000 variable feature computed and used for downstream analyses. Principal component analysis (PCA) has been run and the first 30 principal components (PCs) have been used to find neighbors and clusters at a 0.3 resolution. Finally, the UMAP algorithm has been run with the first 30 PCs, using the “*umap-learn*” algorithm. Seurat objects related to the four datasets have been merged, and the “harmony integration” method has been used to integrate layers. After integration, layers have been joined, and the standard Seurat pipeline has been run to cluster cells and perform all the downstream analyses, using the “*UMAP harmony*” as default reduction.

### Cell population classification and markers

To find markers for each cluster, the “*FindAllMarkers*” function in *Seurat* has been used, with a minimum percentage of expression of 25% and a log2 fold change threshold of 0.25, retrieving both positive and negative markers. Gene ontology analysis on population markers has been performed using “*ClusterProfiler*” (Wu et al., 2021) R package, utilizing all ontologies (GO BP, GO CC, and GO MF), and 10,000 permutations. Results from the functional enrichment analysis, together with the top expressed markers (positive markers) for each cell subpopulation have been used to assign a functional profile to each microglial cell subpopulation.

### Inflammatory score analysis

For the polarization analysis, a customized gene signature was used. The signature was made by manual curation from the literature, using well-known transcription factors and receptors driving macrophage polarization. Specifically, the following genes were used as markers of M1 polarization: *ITGAM, ITGAX, ITGAV, CD36, CD40, CD68, CD80, CD86, CD47, STAT1, STAT3, FCGR2, FCGR3*, and the following genes were used as markers of M2 polarization: *STAT6, ROCK2, TREM2,CD163, PPARG, MRC1, CLEC10A, TGM2, ARG1*. The “*AddModuleScore*” function of the *Seurat* package was then used to compute M1 and M2 scores for each cell. The M1 or M2 identity has been assigned to each cell based on the formula:

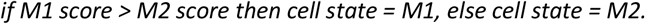

Density plots were drawn using a *ggplot2* customized function, and were edited with *Adobe Illustrator 2024*.

### Pseudobulk between diseases and controls

Pseudobulk analysis was performed using the “*AggregateExpression*” function in Seurat, aggregating cells by disease and patients or harmony clusters. Differential expression was then computed with the “*FindMarkers*” function, using a threshold of adjusted p-value < 0.05 (unless otherwise specified) for determining significantly dysregulated genes for each condition vs ctrl. All control (non-pathological) samples were pooled together and used in each comparison, regardless of their belonging to a specific dataset. Enrichment analysis on GO terms (GO biological processes, cell components, and molecular functions) associated with differentially expressed genes for each pathology was performed with *EnrichR* (Kuleshov et al., 2016) in the R environment, using an adjusted p-value threshold of 0.05 for significant GO terms, and plotting only the top 10 significant terms, based on the adjusted p-value ranking of GO terms. Heatmaps of gene signatures were drawn using the “*pheatmap*” (Kolde, 2019) package. Volcano plots were drawn using the “*EnhancedVolcano*” package. All images were edited with *Adobe Illustrator 2024*.

### Random forest model for disease status classification

The normalized RNA data were extracted from the *Seurat* object, and a subset was created containing only the genes belonging to the neurodegenerative signature (‘AC079142.1’, ‘AC108062.1’, ‘AES’, ‘AL163541.1’, ‘AL392172.2’, ‘AL731577.2’, ‘ANAPC16’, ‘AP001977.1’, ‘ATXN7’, ‘BTBD11’, ‘CERT1’, ‘CRAMP1’, ‘CYRIA’, ‘CYRIB’, ‘DDX39B’, ‘DPH6’, ‘FAM53B’, ‘FP236383.3’, ‘HEXA’, ‘IFI44L’, ‘IFNAR2’, ‘IL10RB’, ‘IRAG2’, ‘LINC00472’, ‘LINC00486’, ‘MARCHF1’, ‘MARCHF3’, ‘MFSD14A’, ‘MRTFB’, ‘MS4A7’, ‘MT-CO3’, ‘MT-ND4’, ‘MTG1’, ‘MUM1’, ‘NDUFA9’, ‘PRKY’, ‘RASGEF1B’, ‘RBM14’, ‘RESF1’, ‘RNASET2’, ‘SEPTIN6’, ‘SGO1-AS1’, ‘SLC26A3’, ‘SNHG14’, ‘SPATA13’, ‘SPECC1L’, ‘UBE2F’, ‘UBE2V1’, ‘UBE2W’, ‘YAE1D1’, ‘ZNF559-ZNF177’, ‘ZNF655’). The model was trained using the “*train*” function of the *caret* package, with the subset data as features and the disease conditions, including controls (CTRL), as labels. The “*random forest*” (*rf*) method was employed for training.

Predictions were generated using the *predict* function, followed by the calculation of the confusion matrix. Feature importance was determined using the *varImp* function. Correct and incorrect predictions were visualized on the UMAP projection by utilizing UMAP coordinates and metadata from the Seurat object, implemented through a customized R script.

ROC curves and AUC values were computed with the *caret* package and plotted with in-house scripts.

The image of the confusion matrix was edited using *GraphPad Prism v7*, and the UMAP images were refined with *Adobe Illustrator 2024*.

### High dimensional weighted gene correlation network analysis

Weighted gene correlation network analysis was performed using the “*hdWGCNA*” package (Morabito et al., 2023). The “fraction” gene selection method was used for setting up the single-cell object, utilizing a threshold of 0.05. Then, the metacell dataset was built grouping by clusters and patients, and the first 6 clusters (from 0 to 5) were used for setting the expression matrix, as no cells were kept by the algorithm for clusters 6, 7, and 8. The standard pipeline provided within the package vignette (https://smorabit.github.io/hdWGCNA/articles/basic_tutorial.html) was then run. For differential module eigengene (DME) analysis between conditions, the pipeline provided in https://smorabit.github.io/hdWGCNA/articles/differential_MEs.html has been used. was used. Enrichment analysis of each module was performed with the “*RunEnrichr*” function provided within the package. Bar plots were drawn with GraphPad (Prism) v7. The network of the top 20 hub genes for each module was retrieved using the “*HubGeneNetworkPlot*” function. The network was transformed into an *igraph* object using the function “*as_data_frame*”. The network was drawn with *Cytoscape v8* (Shannon et al., 2003). All images were edited with *Adobe Illustrator 2024*.

### Statistical analysis and figures

Statistical values in charts are reported as mean ± SEM unless otherwise specified. The statistical significance is defined as Bonferroni adjusted p-value where *p < 0.05; **p < 0.01; ***p < 0.001; ****p < 0.0001. For each type of analysis see the specific Materials and Methods section and Figure captions. Figures were prepared and edited using *Adobe Illustrator 2024* and *GraphPad* (*Prism*) v7 software.

## Results

### Functional heterogeneity of microglial cell population

Publicly available single-nuclei RNA sequencing data were retrieved from the Gene Expression Omnibus (GEO) database. The datasets comprised post-mortem, prefrontal cortex samples from individuals with ALS, AD, PD, FTLD, AGING, and their non-pathological counterparts (Figure 1a). After integrating the datasets (Supplementary Figure 1) and applying quality filtering, 19,537 cells were retained and underwent downstream processing. Clustering analysis revealed the existence of 9 sub-populations (Figure 1b), indicative of the functional heterogeneity of microglial cells.

**Figure 1.**
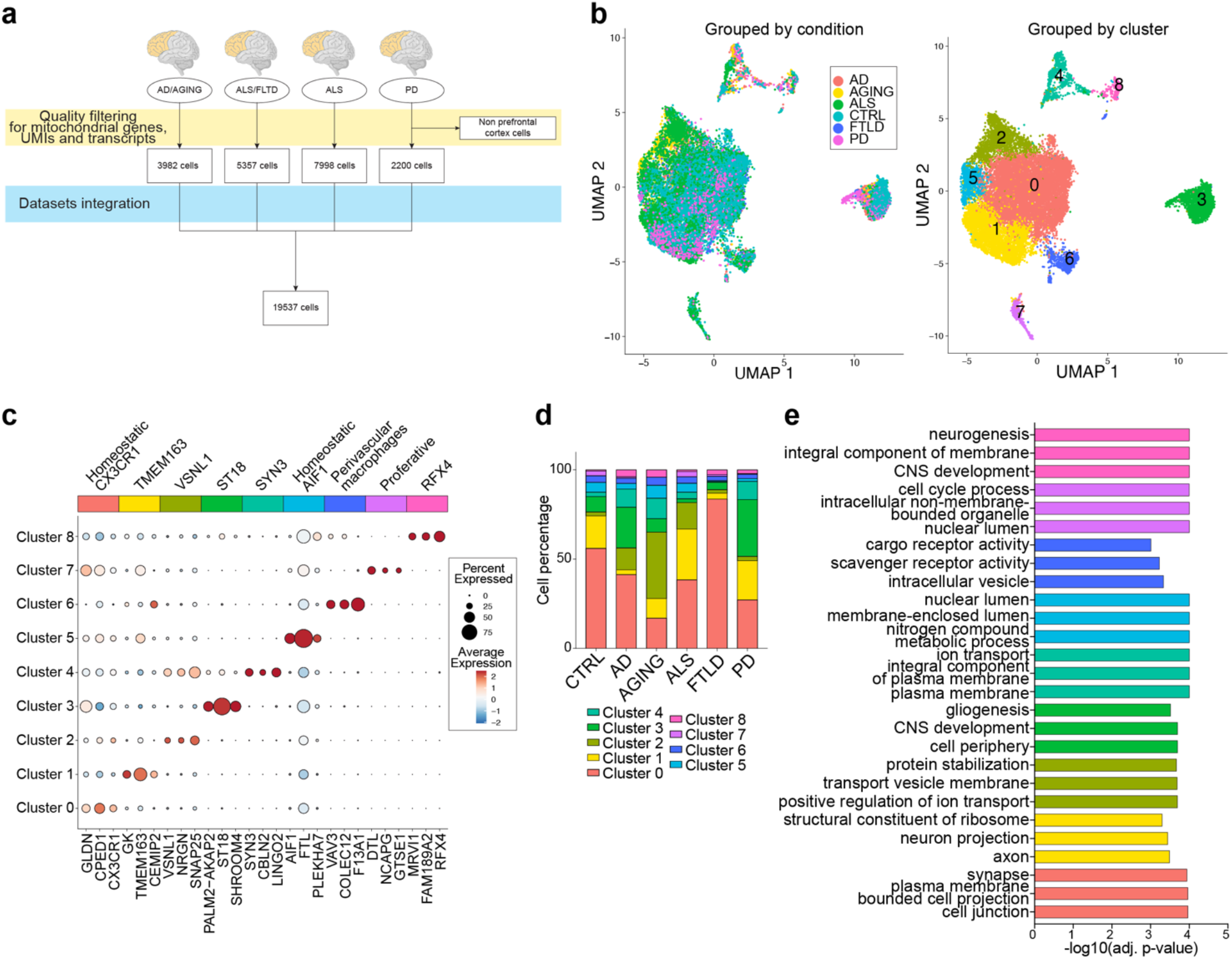
Single-cell analysis of microglia in health and neurodegeneration. (a) Sample stratification chart reporting the number of cells for each dataset and the final number of cells after quality filtering. (b) UMAP projection of the merged dataset showing the overlap between conditions (left panel) and microglial clusters (right panel). (c) Dot plot showing the top 3 markers for each cluster, the percentage of expression, and the average expression of each marker. (d) Stacked bar plot showing the cell count for each cluster grouped by condition. Clusters are colored according to the UMAP cluster colors in panel b. (e) Bar plot showing the top 3 Gene Ontology (GO) biological processes and cellular components derived from the enrichment analysis for each cluster. Bars representing the enrichments are colored according to the clusters in panel b.

The differentially expressed genes (DEGs) specifically marking each cell cluster (Supplementary Table 1) were utilized to elucidate the functional heterogeneity of microglial cells. In particular, genes exhibiting positive values (log 2 fold change > 0) were used as markers to retrieve information about the functional identity of the individual sub-populations (Figure 1c). While several markers were found to be expressed by different clusters to varying extents, an overarching pattern emerged, revealing a distinctive signature for certain sub-populations. For instance, Cluster 0 exhibited expression of well-known microglial markers such as *CPED1* and *CX3CR1*, alongside the oligodendrocyte gene *GLDN*, also detected in clusters 3 and 7. Notably, Cluster 0 uniquely up-regulated microglial markers *P2RY12, FRMD4A*, and *PICALM*, indicative of a homeostatic identity. This cluster was associated with cellular processes related to cell projection and junction (Figure 1e), with cell abundance notably decreasing with aging. Cluster 1 displayed elevated levels of glycerol kinase (*GK*), a microglial marker (*TMEM163*), and *CEMIP2*, which is also expressed in other brain cells. Additionally, it exhibited an adaptive gene signature, including the expression of *HIF1A*. Enrichment analysis revealed associations with processes linked to crosstalk with neuronal populations, and their marked reduction in terms of abundance was also observed in AD and FTLD (Figure 1d). Cluster 2 exhibits elevated levels of *VSNL1*, associated with Alzheimer’s Disease (Groblewska et al., 2015), *NRGN*, a neuron marker, and *SNAP25*. Other markers include mitochondrial genes (MT-genes, *GAPDH*, ATP-genes), and calmodulins (*CALM1, CALM2*). Enrichment analysis revealed associations with processes linked to vesicle and ion transport, with increased cell numbers observed in AD, ALS, and AGING conditions. Cluster 3 is characterized by high expression levels of the gene fusion *PALM2-AKAP2*, the transcription factor *ST18*, which are transcription factors that function as repressors at promoter sites and have been linked in macrophages to the cytokine-mediated signaling pathway and VEGF-A modulation (Maruyama et al., 2020). Furthermore, this cluster expresses high levels of *SHROOM4*, which is involved in cytoskeletal architecture. Notably, cluster 3 is enriched in processes related to gliogenesis and glial cell differentiation, suggesting that this sub-population could serve as precursors to mature glial cells. Intriguingly, cluster 3 cell abundance increased in AD and PD, while a marked decrease was observed in ALS and FTLD. This disparity underscores a potential functional significance of gliogenesis in these diseases. Cluster 4 exhibits high levels of synaptic genes, including *SYN3, CBLN2*, and *LINGO2*, suggesting a potential role for this population in microglia-neurons communication. The abundance of this cell population increased in ALS, AD, PD. Cluster 5 is characterized by the expression of *AIF1, FTL*, and *RPLP2*, markers associated with homeostatic microglia. This cell population is enriched in genes involved in translation, complement activation, and nitrogen metabolic processes, and its abundance is diminished in FTLD and PD. Cluster 6 over-expresses genes encoding cargo and scavenger receptors (*F13A1, MRC1, CD163*), indicative of a perivascular macrophage identity. The abundance of cells in this cluster slightly decreased in FTLD and PD. Consistent with other reports, a cluster of proliferating cells was identified, displaying high expression levels of *MKI67* and other cell cycle-related markers (*TOP2A, NCAPG, KIF18B, POLQ*, among others). Enrichment analysis revealed terms related to the cell cycle and proliferation, with reduced abundance observed in AD, aging, FTLD, and PD. Finally, cluster 8 exhibits high levels of *MRVI1* (*IRAG1*), known to regulate IP3-induced calcium release, *ENTREP1* (*FAM189A2*), which negatively regulates CXCR4 triggering its endocytosis, and transcription factors related to brain system development such as *RFX4* and *EMX2*.

Collectively, these findings suggest that microglia are functionally heterogeneous cell populations, involved in various cellular processes, both in normal physiology and pathology. This heterogeneity is reflected in distinct groups of cells with unique transcriptional profiles. Two clusters showed an homeostatic transcriptional profile, while the others, excluding perivascular macrophages, are more likely identifiable as adaptive microglial populations, each with their own, distinct gene signature. Furthermore, increased or decreased numbers of microglial sub-populations can be observed for distinct disease conditions, which may be a consequence or a pathogenetic mechanism characterizing the different neurodegenerative disorders.

### Pseudobulk analysis reveals a common neurodegeneration gene signature

To assess the existing differences between disease and control samples, a pseudobulk analysis was conducted, aggregating cell transcriptomes by disease and patients, and comparing each disease to the pooled control samples. The pseudobulk analysis revealed hundreds of differentially expressed genes (DEGs), being both up- and down-regulated, in AD, AGING, FTLD and PD conditions (Figure 2a, Supplementary Table 2). No DEGs were found when comparing ALS patients to controls. Consequently, downstream analyses focused on the other disease conditions.

**Figure 2.**
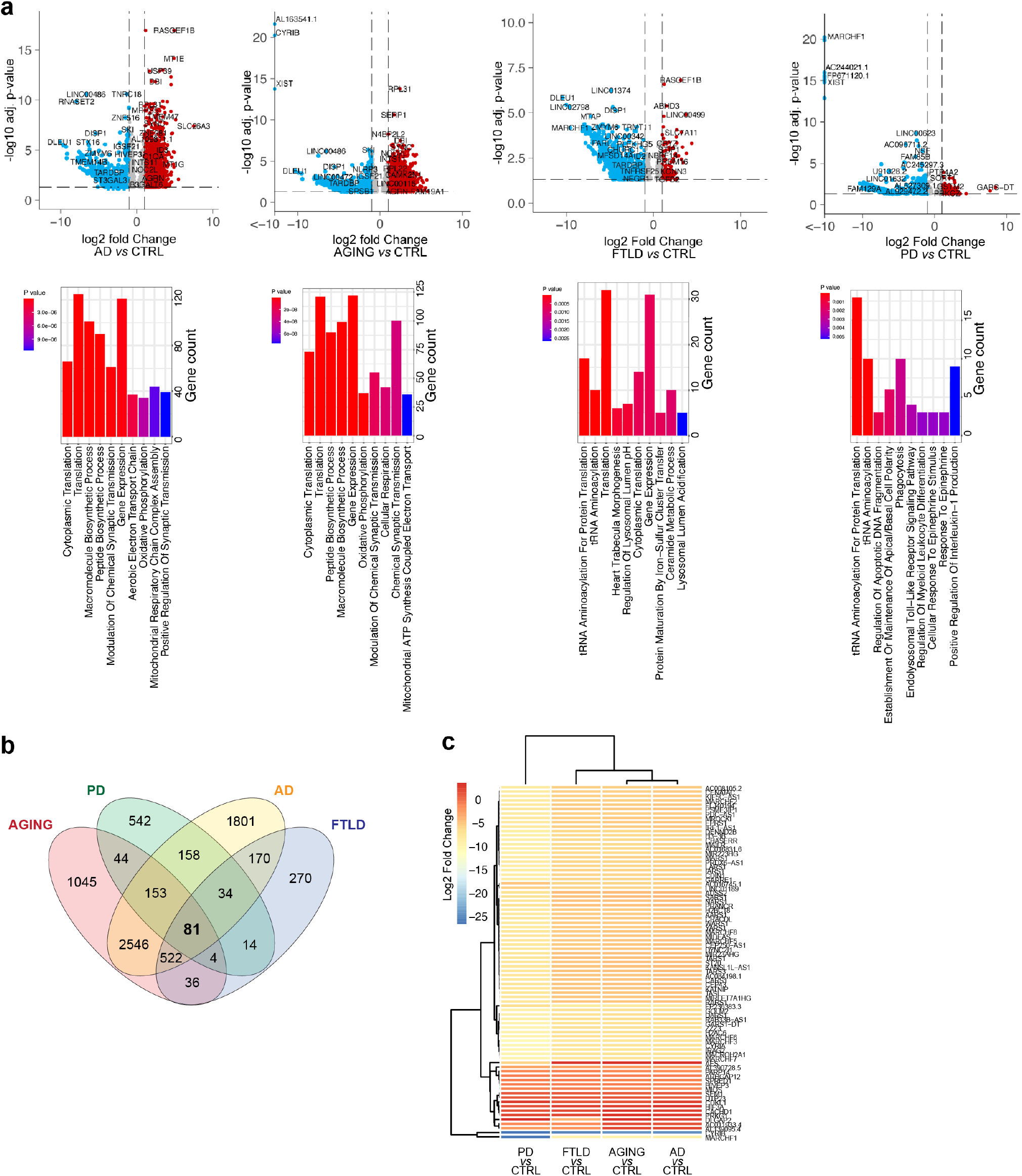
Pseudobulk analysis of AD, aging, ALS, and FTLD vs CTRL. (a) Volcano plots (top panels) showing down-regulated (blue circles) and up-regulated (red circles) genes for each indicated comparison, along with the top 10 enriched GO terms (bottom panels). (b) Venn diagram representing the intersection of DEGs for the reported comparisons. (c) Heatmap of log2 fold changes of the 80 commonly dysregulated genes for the reported comparisons.

In AD samples, DEGs were enriched in Gene Ontology (GO) terms associated with cytoplasmic translation, synapse plasticity, and mitochondria-related metabolic processes (see Supplementary Table 3 for the complete enrichment results). A similar enrichment pattern was observed in the AGING condition, while in FTLD, genes were enriched for lysosome and tRNA aminoacylation. In PD, the top GO terms were associated with the immune response, and, similar to FTLD, to tRNA aminoacylation. These results support common processes underlying different neurodegenerative conditions, as well as distinct and unique cellular events characterizing each disease.

The intersection of common DGEs revealed the presence of 81 genes that were consistently dysregulated across the disease conditions, with the majority showing down-regulation (Figure 1 b, c). These dysregulated genes exhibited agreement in their sign (absolute log2 fold change) and displayed similar expression patterns, suggesting shared transcriptional mechanisms occurring within microglial populations in neurodegenerative conditions.

In particular, the gene signature encompassed several members of the membrane-associated RING-CH-type finger (MARCH) proteins of E3 ubiquitin ligases, which play important roles in immune responses (Lin et al., 2019), genes coding for several aminoacyl-tRNA synthetase, whose mutations have been found in a variety of neurodegenerative and neurological conditions (Sauter et al., 2015), and a number of anti-sense long non-coding RNAs that are yet to be characterized in both health and disease.

Next, cells were aggregated by clusters and disease status to examine differences in gene expression patterns between clusters within each disease condition, potentially revealing cellular heterogeneity and disease-specific effects on different cell types (Supplementary Table 4). Comparing the DEGs in AD, AGING, PD, and FTLD vs CTRL, 52 DEGs were found in common. The gene signature exhibited consistency in directionality and comparable expression levels, as measured by log2 fold change, across AD, AGING, and FTLD. Conversely, a degree of anti-correlation was noted in PD, albeit with limited statistical significance (Supplementary Figure 2). This observation suggests that AD, AGING, and FTLD may share closer transcriptional profiles compared to PD. The neurodegeneration signature comprises long non-coding RNAs, including genes coding for the MARCH protein family (*MARCHF1, MARCHF3*), genes involved in small GTPase activity (*CYRIA, CYRIB, SPATA13*), as well as long non-coding RNAs (*SNHG14, LINC00486, LINC00472*), among others. These genes were among the most down-regulated ones in almost all conditions.

To assess whether this gene signature could classify disease states, it was used to train a random forest model. The resulting model demonstrated its ability to classify cells based on their disease status in the integrated single-cell dataset (Figure 3 a, b) with optimal overall accuracy (accuracy=0.9881, 95% CI=(0.9865, 0.9896) kappa=0.9838), also confirmed by the ROC curves and AUC values (Figure 3 c). This gene signature, encompassing a range of dysregulated genes common across diseases, demonstrated potential as a robust classifier for health and diseased microglial cells. Among the top genes, *SLC26A3, LINC00486* and *MT-CO3* stand out, covering the highest values in the model most important features (Figure 3 d). It has been reported that high levels of *MT-CO3* mRNA have been found in extracellular vesicles from AD patients (Kim et al., 2020), and the gene was also detected high levels in brain microglia from a mouse model of aging (Golomb et al., 2020).

**Figure 3.**
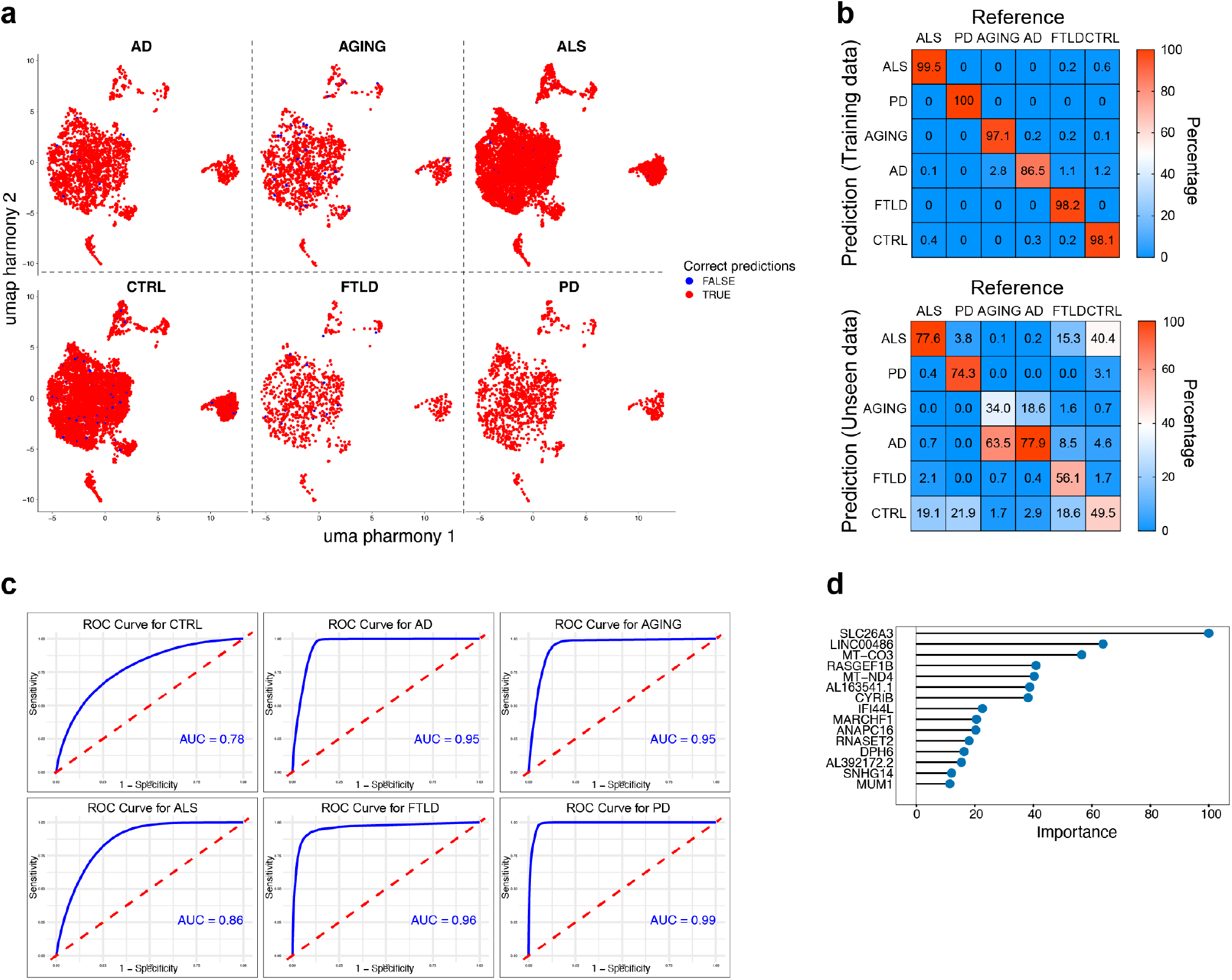
Results of the model build on the neurodegenerative gene signature. (a) UMAP projections split by disease state, showing correct predictions (red labels) and incorrect predictions (blue labels). (b) Confusion matrix of the random forest model, where the number of predicted instances in comparison to references are reported as percentages of the total instances for each disease state. Top panel represent accuracy on the training set, bottom panel shows predictions on new (unseen) data. (c) ROC curves with AUC values (in blue) of the trained model for the prediction accuracy of the reported conditions. (d) Top 15 most important features (genes) derived from the trained model.

### Microglia are not subjected to macrophage-like polarization

The plasticity of myeloid cells, particularly circulating and resident macrophages, has been a subject of debate for many years. The M1/M2 (pro-inflammatory/anti-inflammatory) paradigm has been extensively studied in circulating macrophages, leading to a general agreement regarding the diverse spectrum of polarization states that characterize macrophage activity (Mosser and Edwards, 2008; Sica and Mantovani, 2012). In the context of brain microglia, increasing evidence prompted scientists to classify microglia into different functional states based on their gene expression, in vivo behavior, and morphology under specific experimental conditions or circumstances, rather than adhering strictly to a macrophage-like polarized state (Butovsky and Weiner, 2018; Ransohoff, 2016; Tan et al., 2020). Based on these considerations, and to gain further insights into microglia plasticity in both physiology and disease, an inflammation gene signature (M1/M2-like) was constructed and utilized to generate single-cell scores assigned to each cell. This signature was developed by systematically mining the literature and incorporating well-known pro-inflammatory (*ITGAM, ITGAX, ITGAV, CD36, CD40, CD68, CD80, CD86, CD47, STAT1, STAT3, FCGR2, FCGR3*) and anti-inflammatory (*STAT6, ROCK2, TREM2, CD163, PPARG, MRC1, CLEC10A, TGM2, ARG1*) receptors and transcription factors. The assignment of scores revealed that microglia are not universally in a polarized state (Figure 4 a), with more than half of healthy microglia exhibiting a low pro-/anti-inflammatory score state. This state was also observed in disease conditions, except for aging, where a slight increase in the anti-inflammatory score and the number of cells displaying that inflammatory state was observed (Figure 4 b). Conversely, a marked decrease in both scores was observed in PD. However, upon examining the inflammation score by clusters, it was noted that cluster 6 (perivascular macrophages) contributed to the majority of the anti-inflammatory status of cells (Figure 4 c, d). Considering that the abundance of cluster 6 was decreased in FTLD and PD, and that the anti-inflammatory score was significantly reduced only in PD, it could be speculated that the diminished anti-inflammatory score in PD may be attributed to the depletion or absence of this cell population.

**Figure 4.**
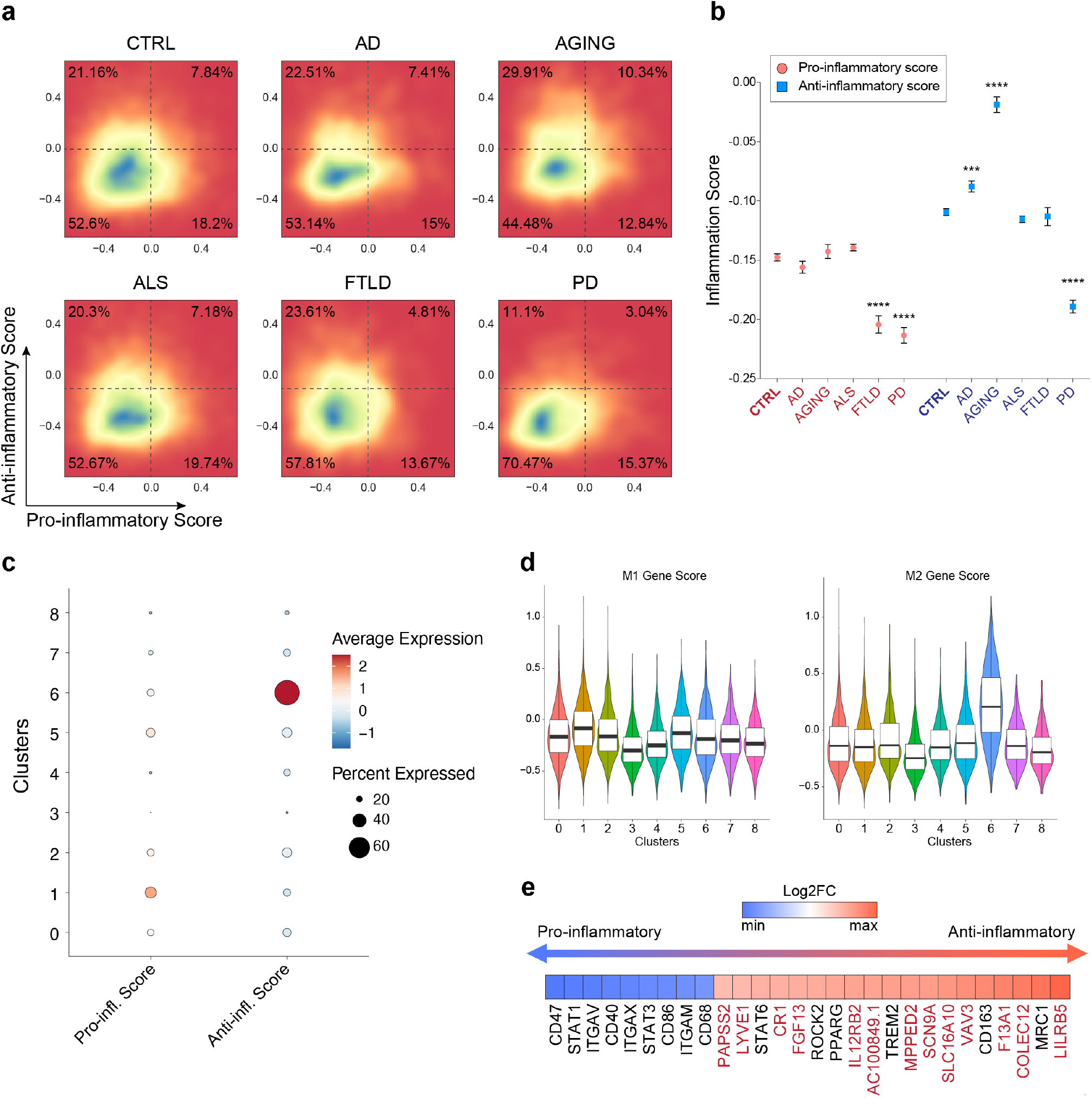
Analysis of the microglial inflammation signature. (a) Density plots illustrating the changes in pro- and anti-inflammatory scores computed using the “AddModuleScore” function for each condition. Percentages of cell numerosity for each quadrant are also depicted. (b) Boxplot presenting the quantification of the inflammation score for each condition. Statistical significance (indicated by stars) pertains to the comparison of each disease condition vs CTRL, conducted separately for pro- and anti-inflammatory scores. (c) Dot plot displaying the expression levels (average expression and percentage) of pro- and anti-inflammatory scores for each cluster. (d) Violin plots with boxplot showing the M1 and M2 gene score distribution across cell clusters. (e) Heatmap reporting the log2 fold change values of the most differentially expressed genes between anti- and pro-inflammatory microglial cells in a blue-white-red scale.

These results underscore that, beside the observation of slight increased or decreased inflammatory states for some brain-resident microglia, this cell population is not classifiable in M1/M2-like polarization states, and that the overall change in polarization/inflammation status observed in certain cells should be mainly attributed to perivascular macrophages.

To improve the list of candidate genes representing the inflammatory signature, a differential gene expression testing has been conducted comparing anti-inflammatory with pro-inflammatory classified microglial cells. The analysis revealed the existence of several differentially expressed genes (Supplementary Table 5) 28 of which with a |log2 fold change| > 1. Among them, 19 were up-regulated in anti-inflammatory-classified cells (anti-inflammatory genes) and 9 were down-regulated in anti-inflammatory cells (pro-inflammatory genes) (Figure 4 e). Some of them were included, as expected, in the gene signature previously used to create the polarization module score and are already known to be associated to a pro-inflammatory or anti-inflammatory-like phenotype, while some others were new. The latter include *LYVE1*, a known macrophage marker implicated in glycosaminoglycan metabolism, *CR1* and *F13A1*, involved in complement and coagulation cascades, with *CR1* being implicated in AD (Brouwers et al., 2012), and *LILRB5*, that act in immune responses, among others. Surprisingly, *IL12RB2*, which is known to trigger a Th1 responses via *IL2* signaling (Liao et al., 2011), was found up-regulated in anti-inflammatory cells.

Overall, the analysis highlights a critical role of perivascular macrophages in influencing the general inflammatory status of brain-resident microglia. Further investigation into the gene expression profiles of microglial subtypes has uncovered new candidate genes associated with inflammatory responses, which could be further studied to better understand the regulation of microglial function in both physiology and disease.

### Weighted gene correlation network analysis identifies differentially expressed networks

To evaluate the presence of gene regulatory networks that may not be fully captured through gene expression measurements alone, a high-dimensional weighted gene correlation network analysis (hdWGCNA) was conducted. This framework enables the construction and analysis of co-expression network data. The analysis identified 8 distinct modules (Figure 5 a). Notably, when assessing module activity within a two-dimensional projection (UMAP), it became evident that certain modules exhibited enrichment within specific microglial subpopulations (Figure 5 b-c). For instance, “Module-1” displayed a strong association with clusters 3, 4, and 8, which represent subpopulations involved in gliogenesis, neurogenesis, and plasma membrane modeling. Intriguingly, this module showed upregulation in aging and PD, while being downregulated in AD (Figure 5 d). Additionally, this module was enriched in genes involved in processes related to the postsynaptic density and neuron projection. This suggests a potential role for this cell population in neuron synapse turnover, which appears to be diminished in AD.

**Figure 5.**
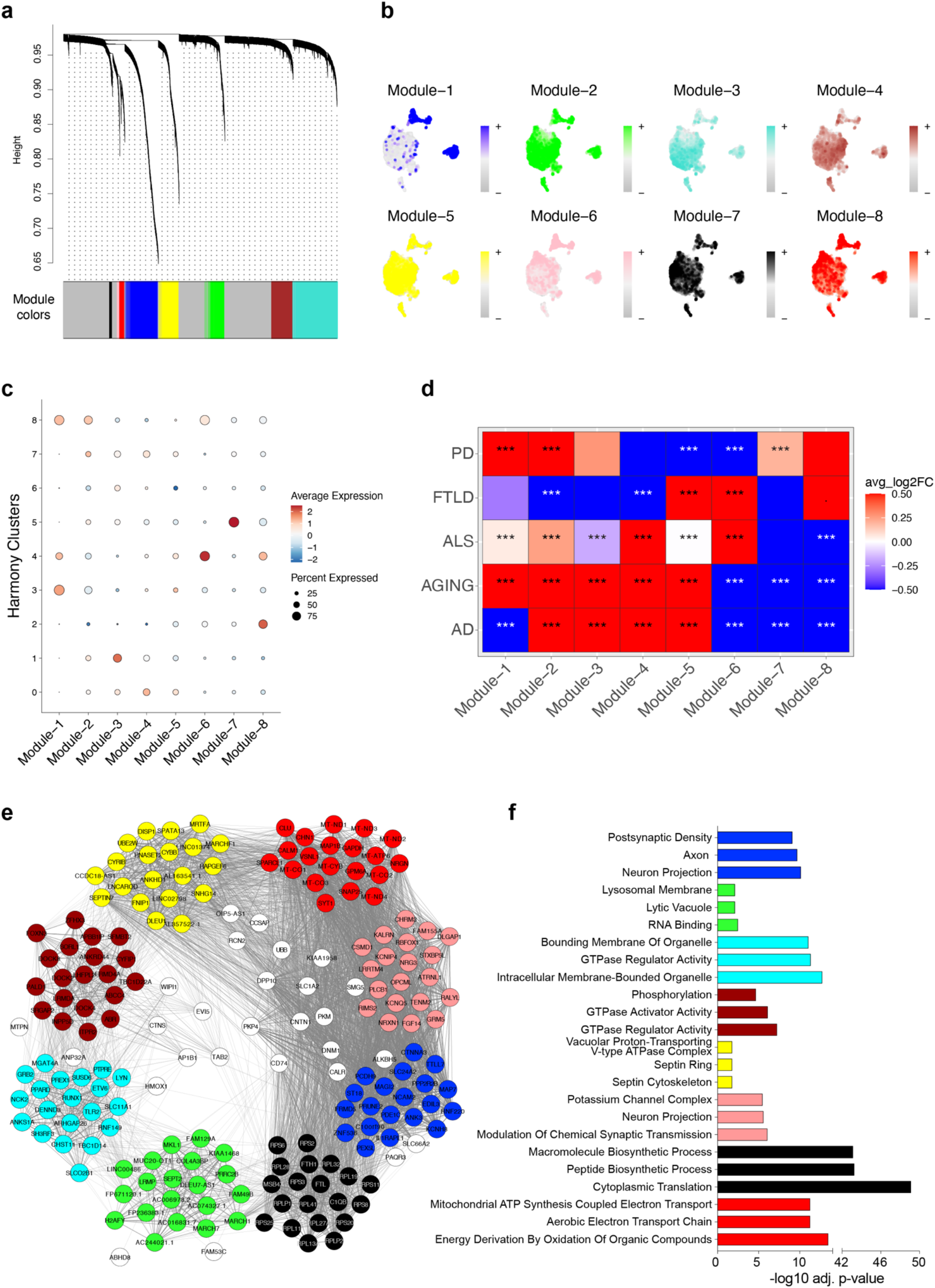
hdWGCNA analysis of the single-nuclei dataset. (a) hdWGCNA dendrogram illustrating the presence of 8 modules (networks). (b) Visualization of module kMEs on the UMAPs. (c) Dot plot displaying the average expression levels and percentage of expression of each module across all clusters. (d) Heatmap presenting the log2 fold change values of all modules for all conditions compared to control samples. (e) Network illustrating the connectivity of the top 20 hub genes for each module, color-coded according to their respective module. (f) Functional enrichment analysis of Gene Ontology Biological Processes and Cell Components for genes within each module. Bars are color-coded according to the module.

Another noteworthy module is “Module-8.” It exhibits significant enrichment in clusters 2 and 8, and it shows marked down-regulation in AD, ALS, and AGING. Within this module, mitochondrial genes rank prominently among the top hub genes, and the module is functionally enriched in mitochondrial metabolic processes. Indeed, numerous studies have underscored the presence of mitochondrial electron transport chain dysfunctions and broader mitochondrial activity impairments in these diseases. Hence, this module offers insights into potential transcriptional dynamics underlying such dysfunctions, providing a set of genes potentially involved in such dysfunctions.

Overall, the enrichment analysis performed on gene regulatory networks partially reflects that obtained from the clustering analysis of microglial sub-populations, confirming dysregulated processes characterizing the distinct clusters, and enriching the picture with additional dysregulated molecular events. A summary of the puzzling picture depicting microglia heterogeneity is provided in figure 6.

**Figure 6.**
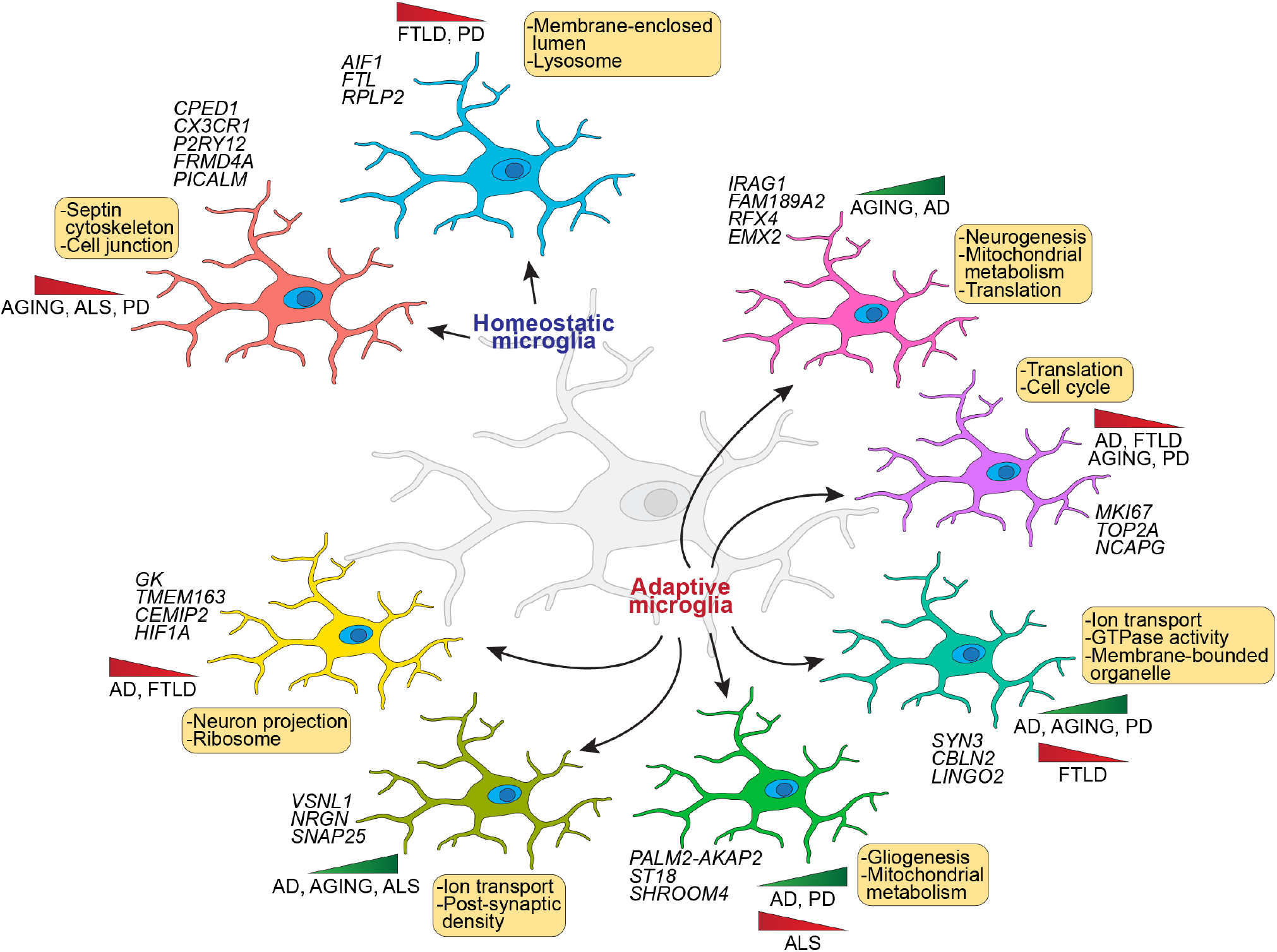
Overall picture of microglia heterogeneity. Picture showing the results obtained from the sub-population identification with clustering analysis and the enriched processes resulted from the weighted gene co-expression network analysis.

## Discussion and conclusions

Microglia, as resident immune cells in the CNS, play diverse and critical roles in maintaining brain homeostasis and responding to pathological conditions. This study utilized and integrated publicly available human single-nuclei RNA sequencing datasets of various neurodegenerative disorders, to unravel the transcriptomic landscape of microglia in health and disease. The findings elucidated the functional heterogeneity of microglial populations, unveiled common neurodegenerative gene signatures, and provided insights into the inflammatory status and gene regulatory networks underlying microglial responses.

The functional heterogeneity of microglial cell populations is highlighted by the clustering analysis, which identified nine distinct clusters exhibiting unique transcriptional profiles and involvement in diverse cellular processes. Notably, certain clusters demonstrated disease-specific alterations in cell numbers, suggesting potential implications for disease pathogenesis. For instance, cluster 3, characterized by genes associated with gliogenesis and glial cell differentiation, showed increased cell numbers in AD and PD, while experiencing a marked decrease in ALS and FTLD. This observation underscores the significance of gliogenesis in specific neurodegenerative contexts and suggests a potential therapeutic target for modulating disease progression.

Furthermore, pseudobulk analysis revealed a shared neurodegenerative gene signature across the analyzed diseases, thus providing insights into common molecular mechanisms underlying neurodegeneration.

The study also investigated the inflammatory status of microglia, challenging the conventional M1/M2 polarization paradigm and revealing a nuanced landscape of pro- and anti-inflammatory states. While a subset of microglia exhibited polarized inflammatory states, a significant proportion maintained a low pro-/anti-inflammatory score, indicative of a non-polarized phenotype. Interestingly, aging was associated with an increase in anti-inflammatory scores, suggesting a potential role of microglia in mitigating inflammation during the aging process. Moreover, the depletion of certain microglial clusters, such as cluster 6 (infiltrating macrophages), in PD was associated with a diminished anti-inflammatory score, highlights the intricate interplay between microglial subpopulations and inflammatory states in disease contexts.

Finally, weighted gene correlation network analysis identified differentially expressed gene networks associated with specific microglial subpopulations and disease states. For instance, dysregulation of mitochondrial metabolic processes, as evidenced by down-regulated modules in AD, ALS, and AGING, underscores the role of mitochondrial dysfunction in neurodegenerative pathologies.

Overall, the analysis unveiled the multifaceted roles of microglia in neurodegenerative disorders and provided a framework for understanding disease-specific alterations in microglial function and gene expression. The limitations of this study include the necessity for further experimental validation of the presented findings, particularly regarding the dysregulation of specific cellular processes occurring within distinct cell subtypes during disease conditions. By elucidating the complex interplay between microglial heterogeneity, inflammatory status, and gene regulatory networks, these results provided the basis for future research aimed at studying targeted strategies to modulate microglial responses and mitigate neurodegenerative disease progression.

## Supplementary material captions

***Supplementary figure 1. Dataset integration***

*UMAP projections showing the overlap between the different datasets before (left panel) and after (right panel) harmony integration*.

***Supplementary figure 2. Gene signature values in the different conditions***

*Pairwise scatterplots showing the comparison of log2 Fold Changes values (disease vs CTRL) for each gene belonging to the signature. Grey shadows represent the confidence intervals. The red dashed lines indicate the curve fitted. R values and p-values are also shown for each comparison*.

***Supplementary Table 1***

*Table reporting the computed markers for each cell subpopulation (cluster). P-values and adjusted p-values, log2 fold changes, clusters, and percentage of cells expressing each marker (pct*.*1 for the considered cluster, pct*.*2 for all the other clusters) are reported*.

***Supplementary Table 2***

*Table reporting the statistics for differentially expressed genes (DEGS) for each condition against all control samples, related to the pseudobulk analysis performed on cells aggregated by disease condition and patients. File sheets are separated by condition*.

***Supplementary Table 3***

*Table reporting the functional enrichment analysis performed on statistically significant genes resulted from the pseudobulk analysis for each disease condition. The data report enrichment statistics related to the enrichment performed on GO Biological Processes (GO BP), GO Cell Component (GO CC) and GO Molecular Function (GO MF) pulled together for each enrichment analysis (each disease vs ctrl)*.

***Supplementary Table 4***

*Table reporting the statistics for differentially expressed genes (DEGS) for each condition against all control samples, related to the pseudobulk analysis performed on cells aggregated by disease condition and harmony clusters. File sheets are separated by condition*.

***Supplementary Table 5***

*Table reporting the statistically significant genes resulted from the differential expression analysis performed on microglial cells classified as anti-inflammatory against microglial cells classified as pro-inflammatory. P-values and adjusted p-values, log2 fold changes, and percentage of cells expressing each gene (pct*.*1 for the anti-inflammatory cells, pct*.*2 for pro-inflammatory cels) are reported*.

## Acknowledgments

The author declares that he is supported by the European Union - NextGenerationEU: National Center for Gene Therapy and Drug based on RNA Technology, CN3 - Spoke 3 (code: CN00000041; PNRR MUR – M4C2 – Action 1.4-Call “Potenziamento strutture di ricerca e di campioni nazionali di R&S”, CUP: B83C22002870006), and that this support exclusively covers his salary.

## Data Availability

Datasets used for this work are available at Gene Expression Omnibus (GEO) repository with the following accession IDs: GSE243292, GSE219281, GSE174332. Parkinson’s disease dataset is available from Zenodo repository at the following link: https://doi.org/10.5281/zenodo.7886802. Code used for this work is available from the corresponding author upon reasonable request.

## References

Bright, F., Werry, E.L., Dobson-Stone, C., Piguet, O., Ittner, L.M., Halliday, G.M., Hodges, J.R., Kiernan, M.C., Loy, C.T., Kassiou, M., et al. (2019). Neuroinflammation in frontotemporal dementia. Nat. Rev. Neurol. 15, 540–555.

Brouwers, N., Van Cauwenberghe, C., Engelborghs, S., Lambert, J.C., Bettens, K., Le Bastard, N., Pasquier, F., Montoya, A.G., Peeters, K., Mattheijssens, M., et al. (2012). Alzheimer risk associated with a copy number variation in the complement receptor 1 increasing C3b/C4b binding sites. Mol. Psychiatry 17, 223–233.

Butovsky, O., and Weiner, H.L. (2018). Microglial signatures and their role in health and disease. Nat. Rev. Neurosci. 19, 622–635.

Crapser, J.D., Arreola, M.A., Tsourmas, K.I., and Green, K.N. (2021). Microglia as hackers of the matrix: sculpting synapses and the extracellular space. Cell. Mol. Immunol. 18, 2472–2488.

Dehestani, M., Kozareva, V., Blauwendraat, C., Fraenkel, E., Gasser, T., and Bansal, V. (2023). Transcriptomic changes in oligodendrocytes and precursor cells predicts clinical outcomes of Parkinson’s disease. BioRxiv.

Gao, C., Jiang, J., Tan, Y., and Chen, S. (2023). Microglia in neurodegenerative diseases: mechanism and potential therapeutic targets (Springer US).

Golomb, S.M., Guldner, I.H., Zhao, A., Wang, Q., Palakurthi, B., Aleksandrovic, E.A., Lopez, J.A., Lee, S.W., Yang, K., and Zhang, S. (2020). Multi-modal Single-Cell Analysis Reveals Brain Immune Landscape Plasticity during Aging and Gut Microbiota Dysbiosis. Cell Rep. 33, 108438.

Groblewska, M., Muszyński, P., Wojtulewska-Supron, A., Kulczyńska-Przybik, A., and Mroczko, B. (2015). The Role of Visinin-Like Protein-1 in the Pathophysiology of Alzheimer’s Disease. J. Alzheimer’s Dis. 47, 17–32.

Hao, Y., Stuart, T., Kowalski, M.H., Choudhary, S., Hoffman, P., Hartman, A., Srivastava, A., Molla, G., Madad, S., Fernandez-Granda, C., et al. (2024). Dictionary learning for integrative, multimodal and scalable single-cell analysis. Nat. Biotechnol. 42, 293–304.

Hickman, S., Izzy, S., Sen, P., Morsett, L., and El Khoury, J. (2018). Microglia in neurodegeneration. Nat. Neurosci. 21, 1359–1369.

Hu, J., Chen, Q., Zhu, H., Hou, L., Liu, W., Yang, Q., Shen, H., Chai, G., Zhang, B., Chen, S., et al. (2023). Microglial Piezo1 senses Aβ fibril stiffness to restrict Alzheimer’s disease. Neuron 111, 15–29.e8.

Joers, V., Mulas, G., and Carta, A.R. (2017). Microglial phenotypes in Parkinson’s disease and animal models of the disease. Prog. Neurobiol. 57–75.

Kent, S.A., and Miron, V.E. (2024). Microglia regulation of central nervous system myelin health and regeneration. Nat. Rev. Immunol. 24, 49–63.

Kim, K.M., Meng, Q., Perez de Acha, O., Mustapic, M., Cheng, A., Eren, E., Kundu, G., Piao, Y., Munk, R., Wood, W.H., et al. (2020). Mitochondrial RNA in Alzheimer’s Disease Circulating Extracellular Vesicles. Front. Cell Dev. Biol. 8, 1–14.

Kolde, R. (2019). Pheatmap: pretty heatmaps.

Kuleshov, M. V., Jones, M.R., Rouillard, A.D., Fernandez, N.F., Duan, Q., Wang, Z., Koplev, S., Jenkins, S.L., Jagodnik, K.M., Lachmann, A., et al. (2016). Enrichr: a comprehensive gene set enrichment analysis web server 2016 update. Nucleic Acids Res. 44, W90.

Leng, F., and Edison, P. (2021). Neuroinflammation and microglial activation in Alzheimer disease: where do we go from here? Nat. Rev. Neurol. 17, 157–172.

Li, Q., and Barres, B.A. (2018). Microglia and macrophages in brain homeostasis and disease. Nat. Rev. Immunol. 18, 225–242.

Liao, W., Lin, J.-X., Wang, L., Li, P., and Leonard, W.J. (2011). Cytokine receptor modulation by interleukin-2 broadly regulates T helper cell lineage differentiation. Nat. Immunol. 12, 551–559.

Lin, H., Li, S., and Shu, H.B. (2019). The Membrane-Associated MARCH E3 Ligase Family: Emerging Roles in Immune Regulation. Front. Immunol. 10, 1751.

Lloyd, A.F., and Miron, V.E. (2019). The pro-remyelination properties of microglia in the central nervous system. Nat. Rev. Neurol. 15, 447–458.

Maruyama, K., Kidoya, H., Takemura, N., Sugisawa, E., Takeuchi, O., Kondo, T., Eid, M.M.A., Tanaka, H., Martino, M.M., Takakura, N., et al. (2020). Zinc Finger Protein St18 Protects against Septic Death by Inhibiting VEGF-A from Macrophages. Cell Rep. 32.

Masuda, T., Sankowski, R., Staszewski, O., and Prinz, M. (2020). Microglia Heterogeneity in the Single-Cell Era. Cell Rep. 30, 1271–1281.

Morabito, S., Reese, F., Rahimzadeh, N., Miyoshi, E., and Swarup, V. (2023). hdWGCNA identifies co-expression networks in high-dimensional transcriptomics data. Cell Reports Methods 3.

Mosser, D.M., and Edwards, J.P. (2008). Exploring the full spectrum of macrophage activation. Nat. Rev. Immunol. 8, 958–969.

Niraula, A., Sheridan, J.F., and Godbout, J.P. (2017). Microglia Priming with Aging and Stress. Neuropsychopharmacology 42, 318–333.

Perry, V.H., and Holmes, C. (2014). Microglial priming in neurodegenerative disease. Nat. Rev. Neurol. 10, 217–224.

Ransohoff, R.M. (2016). A polarizing question: Do M1 and M2 microglia exist. Nat. Neurosci. 19, 987–991.

Ronzano, R., Roux, T., Thetiot, M., Aigrot, M.S., Richard, L., Lejeune, F.X., Mazuir, E., Vallat, J.M., Lubetzki, C., and Desmazières, A. (2021). Microglia-neuron interaction at nodes of Ranvier depends on neuronal activity through potassium release and contributes to remyelination. Nat. Commun. 12.

Salter, M.W., and Stevens, B. (2017). Microglia emerge as central players in brain disease. Nat. Med. 23, 1018–1027.

Sauter, C., Lorber, B., Gaudry, A., Karim, L., Schwenzer, H., Wien, F., Roblin, P., Florentz, C., and Sissler, M. (2015).Neurodegenerative disease-associated mutants of a human mitochondrial aminoacyl-tRNA synthetase present individual molecular signatures. Sci. Rep. 5, 1–13.

Shannon, P., Markiel, A., Ozier, O., Baliga, N.S., Wang, J.T., Ramage, D., Amin, N., Schwikowski, B., and Ideker, T. (2003). Cytoscape: A Software Environment for Integrated Models of Biomolecular Interaction Networks. Genome Res. 13, 2498–2504.

Sica, A., and Mantovani, A. (2012). Macrophage plasticity and polarization: in vivo veritas. J. Clin. Invest. 122, 787–795.

Tan, Y.L., Yuan, Y., and Tian, L. (2020). Microglial regional heterogeneity and its role in the brain. Mol. Psychiatry 25, 351–367.

Touil, H., Li, R., Zuroff, L., Moore, C.S., Healy, L., Cignarella, F., Piccio, L., Ludwin, S., Prat, A., Gommerman, J., et al. (2023). Cross-talk between B cells, microglia and macrophages, and implications to central nervous system compartmentalized inflammation and progressive multiple sclerosis. EBioMedicine 96, 104789.

Vahsen, B.F., Gray, E., Thompson, A.G., Ansorge, O., Anthony, D.C., Cowley, S.A., Talbot, K., and Turner, M.R. (2021). Non-neuronal cells in amyotrophic lateral sclerosis — from pathogenesis to biomarkers. Nat. Rev. Neurol. 17, 333–348.

Wu, T., Hu, E., Xu, S., Chen, M., Guo, P., Dai, Z., Feng, T., Zhou, L., Tang, W., Zhan, L., et al. (2021). clusterProfiler 4.0: A universal enrichment tool for interpreting omics data. Innovation 2, 100141.

