## Supplementary figures and images for "Unveiling microglial heterogeneity from single-cell transcriptomics in neurodegenerative diseases"

### Supplementary Figure 1

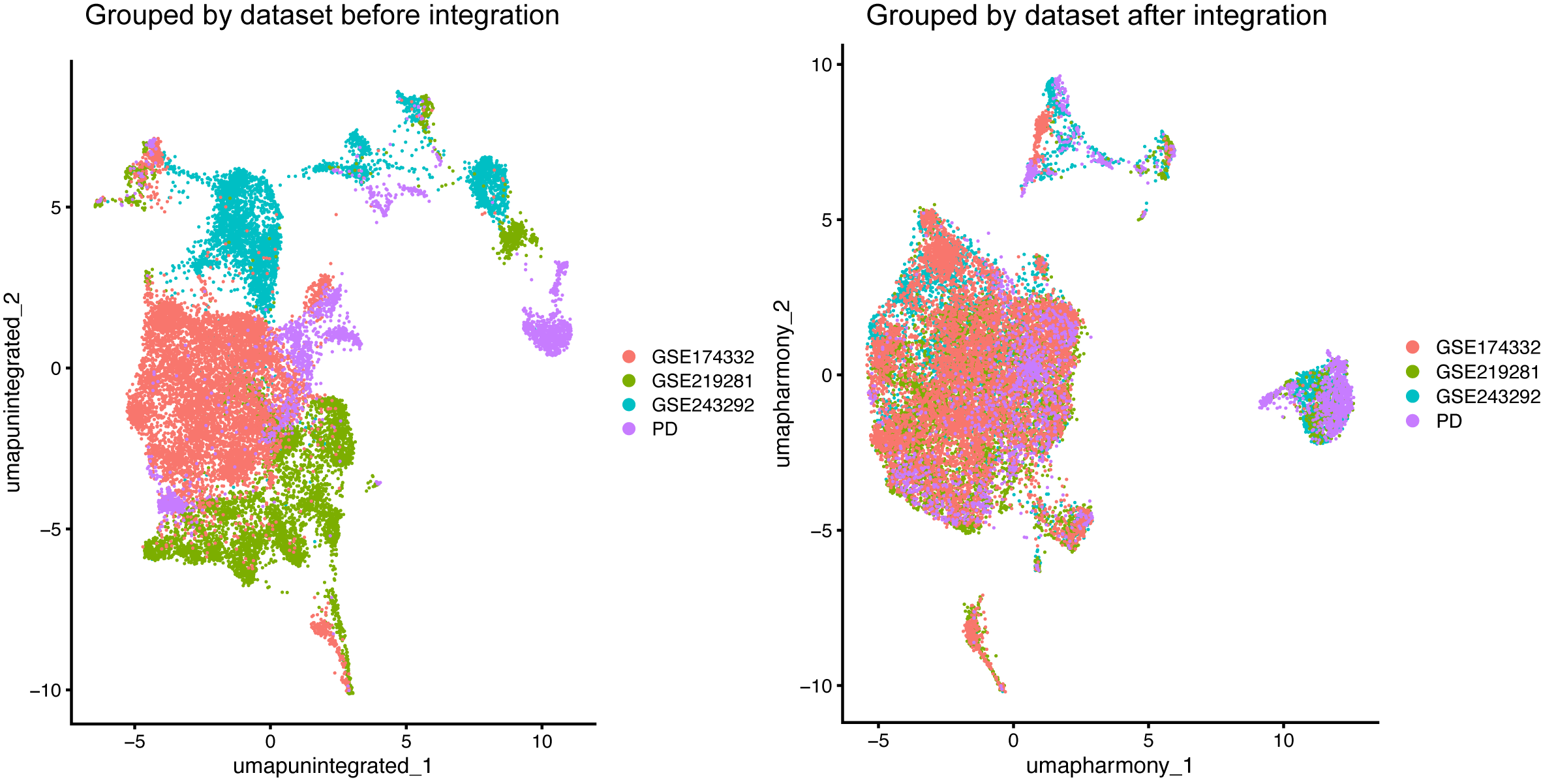

### Supplementary Figure 2

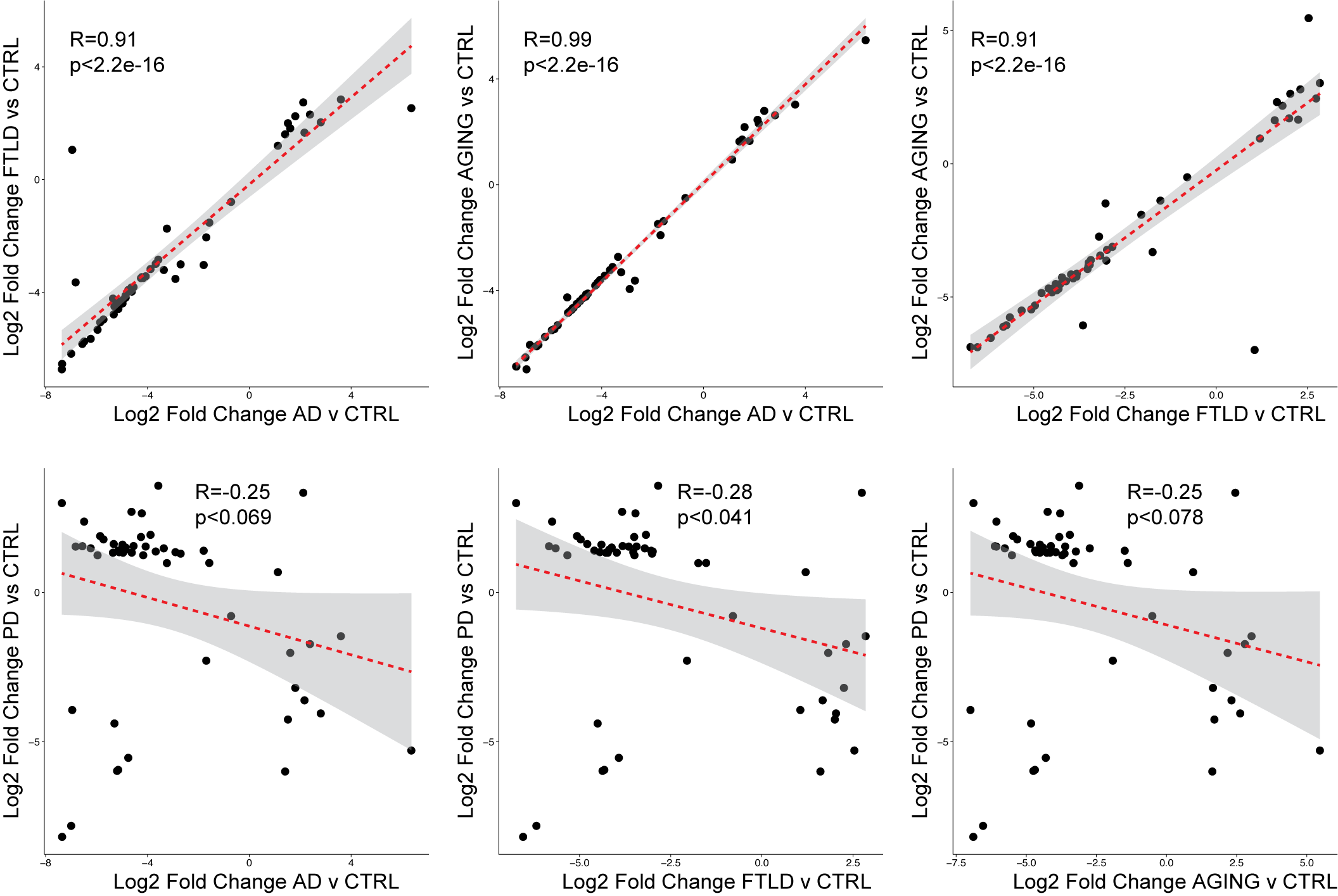
